# Epigenetic regulation of alveolar macrophage homeostasis by histone deacetylase 1 and 2

**DOI:** 10.1101/2024.05.06.590855

**Authors:** Wenlong Lai, Maocai Luo, Yuanhao Wang, Xiaoguang Li, Zhimin He, Tao Wu, Li Wu

**Author notes:** These authors contributed equally (Wenlong Lai, Maocai Luo). Correspondence (T.W.), (L.W.).

## Abstract

Alveolar macrophages (AMs) are the primary lung-resident macrophages that play a pivotal role in pathogen clearance and surfactant homeostasis. Although the molecular regulation of AM development at the transcriptional level has been more widely studied, the epigenetic mechanisms that maintain AM homeostasis remain incompletely understood. Here, we demonstrate that histone deacetylases HDAC1 and HDAC2 play essential roles in preserving AM integrity. Dual deletion of HDAC1 and HDAC2 resulted in severe defect in AM development, leading to pulmonary alveolar proteinosis (PAP) and increased susceptibility to influenza infection in mice. Mechanistically, HDAC1 and HDAC2 act through their deacetylase activity to suppress the expression of pro-apoptotic protein Bim, thereby enhancing cell survival. Furthermore, HDAC1 and HDAC2 form a functional complex with the histone demethylase LSD1 (KDM1A) to co-regulate a transcriptional program governing AM differentiation. Our findings identify HDAC1 and HDAC2 as central epigenetic regulators essential for alveolar macrophage homeostasis.

## Introduction

Alveolar macrophages (AMs) are the major macrophage population residing on the luminal surface of the alveoli. As key immune sentinels in the lung, AMs maintain pulmonary homeostasis by clearing incoming pathogens and airborne pollutants^1^. Their dysfunction causes pulmonary surfactant accumulation, leading to pulmonary alveolar proteinosis (PAP)^2^. AMs primarily originate from fetal monocytes that colonize the lungs before birth and achieve a stable phenotype postnatally^3–6^. Under steady-state conditions, AMs self-maintain through local proliferation without input from bone marrow (BM) precursors^7^. Their identity is heavily shaped by the alveolar microenvironment, particularly GM-CSF secreted by alveolar epithelial cells, which guides their differentiation and maturation^3,5^. Impaired GM-CSF signaling (e.g., in *Csf2*^−/−^, *Csf2rb*^−/−^, or *Csf2ra*^−/−^ mice) leads to AM deficiency^8,9^ and surfactant accumulation^8–12^. GM-CSF drives expression of peroxisome proliferator-activated receptor-gamma (PPAR-γ), a key regulator of lipid metabolism that defines AM identity^5^. Other crucial factors, including mTOR^13,14^, PU.1^15^, C/EBPβ^16^ and STAT5^17^, are also essential for AM differentiation and function under steady-state conditions. Additionally, autocrine TGF-β signaling enhances PPAR-γ expression and promotes AM homeostasis^18^.

Although the roles of cytokines and transcription factors in regulating AM homeostasis are increasingly understood, the epigenetic mechanisms involved remain poorly understood. Histone acetylation plays a central role in transcriptional regulation, which is dynamically regulated by histone acetyltransferases (HATs) and deacetylases (HDACs)^19^. HDACs remove acetyl groups from histones, leading to chromatin condensation and transcriptional repression^20^. Mammalian genomes encode 18 HDACs, which are categorized into four classes based on their structure, enzymatic mechanism and intracellular localization. HDAC1 and HDAC2 belong to class I HDACs, and have a wide tissue distribution and share ∼83% sequence similarity^21^. Both HDAC1 and HDAC2 are critical for immune cell development and function. Their inactivation impairs hematopoiesis^22–24^, lymphopoiesis^25–27^, microglia maintenance^28^, and interferon production by macrophages^29,30^. Notably, in human chronic obstructive pulmonary disease (COPD), reduced HDAC2 expression in AMs is associated with disease severity^31,32^, suggesting a vital role for epigenetic regulation.

In this study, we utilized a genetic approach to conditionally delete both *Hdac1* and *Hdac2* in AMs. Our findings reveal that HDAC1 and HDAC2 are indispensable and exhibit functional redundancy in maintaining AM homeostasis. Deficiency in either *Hdac1* or *Hdac2* alone did not significantly affect AM homeostasis, however, combined deletion resulted in profound defects in AM development, culminating in PAP and significantly enhanced susceptibility to viral infection in mice. We first confirmed that loss of HDAC1 and HDAC2 did not significantly affect GM-CSF signaling or PPARγ expression. We next demonstrated that the deacetylase activity of HDAC1 and HDAC2 was required to repress the expression of pro-apoptotic *Bim*, thereby promoting AM survival. Furthermore, we found that HDAC1 and HDAC2 interacted with the histone demethylase LSD1, enabling coordinated regulation of a transcriptional program essential for AM differentiation. Taken together, these findings reveal HDAC1 and HDAC2 as master epigenetic regulators for AM homeostasis and function through coordinating transcriptional programs governing cell survival and differentiation.

## Results

### Deficiency of HDAC1 and HDAC2 impairs AM homeostasis

To investigate the role of *Hdac1* and *Hdac2* in vivo, we generated mice with conditional deletion of these genes in CD11c^+^ cells by crossing *Hdac1*^fl/fl^ and *Hdac2*^fl/fl^ mice to the *Itgax*-Cre line (Fig. S1A). Efficient deletion (>90%) was confirmed by qPCR in AMs isolated from both *Itgax*^Cre^*Hdac1*^fl/fl^ and *Itgax*^Cre^*Hdac2*^fl/fl^ mice (Fig. S1B). Due to potential functional redundancy between HDAC1 and HDAC2, we also generated *Itgax*^Cre^*Hdac1*^fl/fl^*Hdac2*^fl/fl^ mice (referred to as *Itgax*^Cre^-DKO). AMs were identified as CD45^+^CD11c^+^Siglec-F^+^CD11b^−^F4/80^+^CD64^+^population with high autofluorescence. Single deletion of either *Hdac1* or *Hdac2* did not affect AM numbers or proportions, whereas dual deficiency resulted in a near-complete loss of AMs in *Itgax*^Cre^-DKO mice (Fig. 1A-D). The remaining CD11c^+^ cells in mutants showed reduced autofluorescence and CD64 expression (Fig. S1C), consistent with AM loss. Reductions in CD11b^+^ and CD103^+^ dendritic cell (DC) subsets were also observed (Fig. S2A-D), suggesting broader effects of *Hdac1* and *Hdac2* deletion within CD11c^+^ compartments. These results indicate that HDAC1 and HDAC2 are collectively essential for AM development.

**Figure 1.**
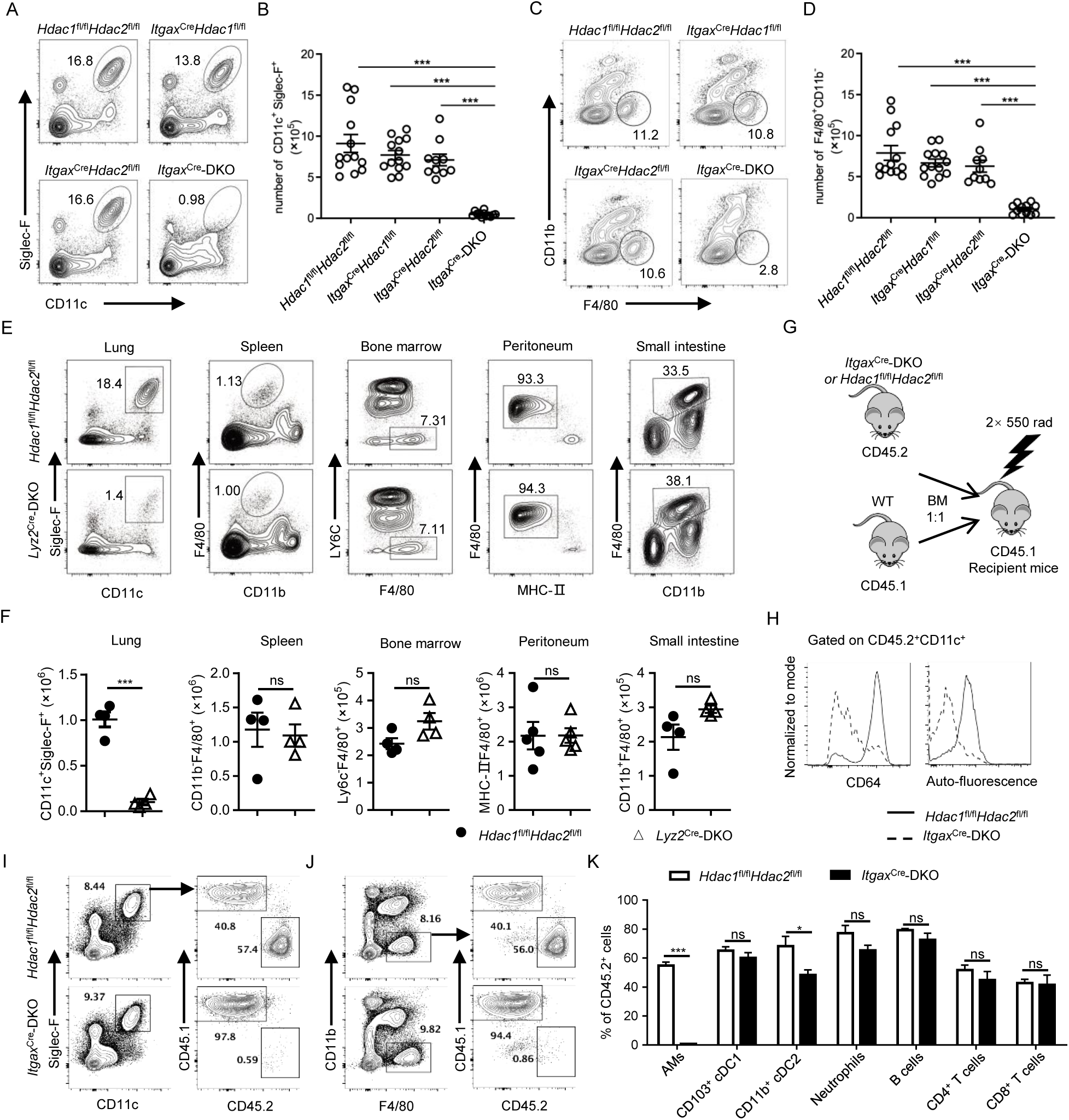
Essential roles of HDAC1 and HDAC2 in AM homeostasis. (A-D). Representative flow cytometry plots (A and C) and cell numbers (B and D) of AMs with different gating strategies: Siglec-F^+^CD11c^+^ (A and B) or F4/80^+^CD11b^−^ (C and D) gated on CD45^+^ cells, (n=10-13). Data were pooled from four independent experiments. (E). Representative flow cytometry plots of AM (gated on CD45^+^), and macrophages in spleen, bone marrow (BM, gated on CD11b^+^Ly6G^−^Siglec-F^−^), peritoneum (gated on CD45^+^CD11b^+^) and small intestine (gated on CD45^+^MHC-II^+^). (F) Cell numbers of indicated tissue macrophages in (E) (n=4-5). Data were representative of two independent experiments. (G). Schematic for the generation of mixed bone marrow chimera. (H). Expression of CD64 and autofluorescence of cells gated on CD45.2^+^CD11c^+^ cells. (I, J). Identification of AMs (SiglecF^+^CD11c^+^ or F4/80^+^CD11b^−^) derived from the CD45.2^+^ donor cells in the lungs of chimeric mice. (K). Frequency of CD45.2^+^ cells among neutrophils, DC subsets, AMs, T cells and B cells in chimeric mice (n=4-5). Data were representative of three independent experiments. Data are shown as mean ± SEM. Statistical significance was determined by one-way ANOVA (B, D) or unpaired two-tailed Student’s *t* test (F, K). *, *p* < 0.05; **, *p* < 0.01; ***, *p* < 0.001.

To further dissect the tissue-specific roles of HDAC1 and HDAC2, we generated myeloid-specific deficient mice by crossing *Hdac1*^fl/fl^*Hdac2*^fl/fl^ mice with *Lyz2-*Cre mice (referred to as *Lyz2*^Cre^-DKO). Consistent with the findings in *Itgax*^Cre^-DKO mice, *Lyz2*^Cre^-DKO mice also exhibited a severe deficiency of AMs in the lungs (Fig. 1E, F). In contrast, macrophages across lymphoid and non-lymphoid tissues, including the spleen, BM, peritoneum, and small intestine, were present at normal frequencies and numbers compared to wild-type controls (Fig. 1E, F). To assess Cre activity across these populations, we introduced a Cre-dependent RFP reporter into the *Lyz2*^Cre^ mice^33^. Cre-mediated recombination efficiency was lower in BM macrophages (∼50% RFP□), but highly efficient in lung, splenic, peritoneal, and intestinal macrophages (75–90% RFP□; Fig. S1D), highlighting tissue-speecific Cre activity. Consistently, qPCR analysis confirmed ∼90% deletion efficiency of *Hdac1* and *Hdac2* in AMs, but less than 50% in splenic, BM and intestinal macrophages (Fig. S1E, F). Collectively, these results demonstrate that HDAC1 and HDAC2 are essential for alveolar macrophage homeostasis.

To determine whether HDAC1 and HDAC2 are intrinsically required for the BM-derived replenishment of AMs, we generated competitive bone marrow chimeras by transplanting a 1:1 mixture of CD45.1^+^ wild-type (WT) BM cells and CD45.2^+^ BM cells from either *Itgax*^Cre^-DKO or control *Hdac1*^fl/fl^*Hdac2*^fl/fl^ mice into lethally irradiated CD45.1^+^ WT recipients (Fig. 1G). Eight weeks post-transplantation, recipient lungs were analyzed by flow cytometry. The CD45.2^+^CD11c^+^ cells derived from *Itgax*^Cre^-DKO BM cells exhibited markedly reduced autofluorescence and CD64 expression compared to controls (Fig. 1H). AMs in mice receiving WT and control *Hdac1*^fl/fl^*Hdac2*^fl/fl^ BM cells showed approximately a 1:1 reconstitution ratio. In contrast, fewer than 1% of AMs in recipient mice were derived from *Itgax*^Cre^-DKO BM cells (Fig. 1I-K). While CD45.2^+^CD103^+^ cDC1 were reconstituted at comparable ratios between groups, CD11b^+^ cDC2 derived from *Itgax*^Cre^-DKO BM showed impaired reconstitution (Fig. 1K), suggesting a broader cell-intrinsic requirement of HDAC1 and HDAC2 in CD11c^+^ lineage cells, including CD11b^+^ cDC2.

Taken together, these results demonstrate that HDAC1 and HDAC2 are cell-intrinsically required for BM-derived contribution to alveolar macrophage maintenance.

### HDAC1 and HDAC2 deficiency in AMs impairs surfactant clearance and antiviral response

AMs are central to pulmonary homeostasis by clearing surfactant secreted by alveolar type II epithelial cells. Deficiency of AMs leads to PAP. To evaluate pulmonary pathology in *Hdac1* and *Hdac2* double deficient mice, we analyzed *Itgax*^Cre^-DKO mice and observed a marked accumulation of total protein levels in bronchoalveolar lavage fluid (BALF), accompanied by intense periodic acid–Schiff (PAS) staining (Fig. 2A-C). Immunoblotting further revealed elevated levels of surfactant proteins in lung tissues (Fig. 2D). Moreover, increased neutrophil counts in BALF (Fig. S3A, B) and elevated levels of pro-inflammatory cytokines (Fig. S3C) were detected, indicating that AM deficiency induces low-grade pulmonary inflammation. Collectively, these findings indicate that loss of *Hdac1* and *Hdac2* in AMs leads to spontaneous development of a PAP-like phenotype.

**Figure 2.**
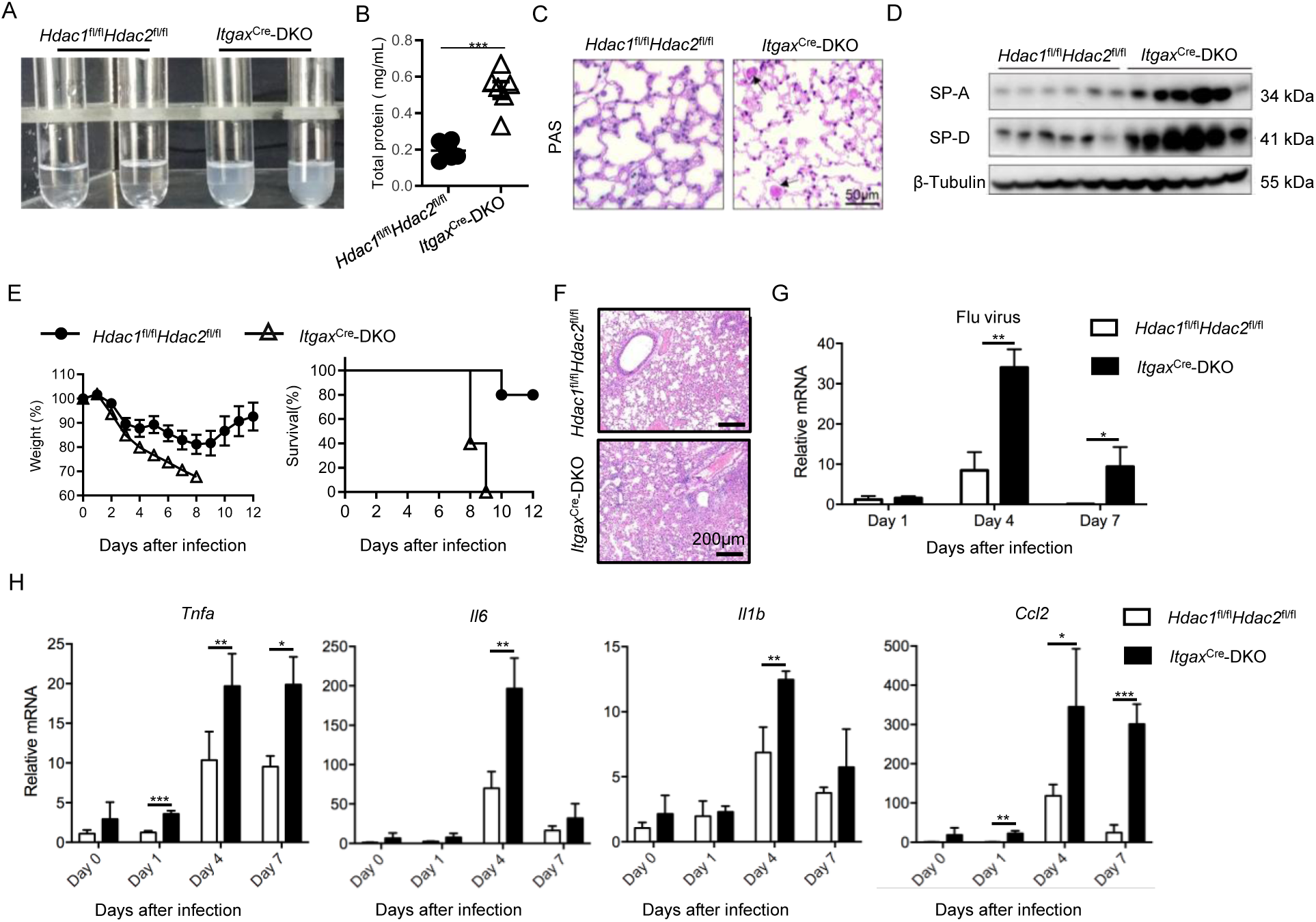
Loss of AMs impairs surfactant clearance and antiviral response in *Hdac1* and *Hdac2* double-deficient mice. (A). Representative images for BALF from mice of indicated genotypes. (B). Quantification of total protein concentration in BALF (n=6). (C). PAS staining of lung tissues from indicated mice at 10 weeks of age. Arrows indicate protein accumulation in the lung. Images were representative of three mice. (D). Western blot analysis of SP-A and SP-D in lung lysates (n=6 mice). (E). Body weight changes and survival curves of PR8-infected mice (500 TCID_50_, n=5). Data were representative of two independent experiments. (F). H&E staining of lung tissue from the indicated mice on day 7 after PR8 infection. Images were representative of three mice. (G-H). Relative mRNA levels of influenza virus (H)*, Tnf*, *Il6, Il1b* and *Ccl2* (I) in lung tissue after PR8 infection (500TCID_50_), normalized to *Actb* expression (n=3). Data were shown as mean ± SEM. Statistical significance was determined by unpaired two-tailed Student’s *t* test. *, *p* < 0.05; **, *p* < 0.01; ***, *p* < 0.001.

Beyond their homeostatic functions, AMs are key effector cells in the innate immune defense against respiratory viral infections. Previous depletion and genetic models have demonstrated a protective role of AMs in limiting influenza virus-induced lethal inflammation^34^. To investigate whether HDAC1 and HDAC2 are required for this function, *Itgax*^Cre^*-*DKO mice and control *Hdac1*^fl/fl^*Hdac2*^fl/fl^ mice were infected intranasally with the PR8 strain of influenza virus, a widely used mouse model of influenza infection. Following virus challenge (500 TCID_50_ per mouse), all *Itgax*^Cre^*-*DKO mice succumbed between day 8 and 10 post-infection, whereas control mice exhibited markedly improved survival (Fig. 2E). *Itgax*^Cre^*-*DKO mice also showed more pronounced weight loss, while control mice began to recover and regain weight around day 8 (Fig. 2E). Histopathological analysis revealed severe pulmonary edema and neutrophil infiltration in *Itgax*^Cre^*-*DKO lungs compared with controls (Fig. 2F). Furthermore, *Itgax*^Cre^*-*DKO mice exhibited significantly higher viral titers at the peak of infection (day 4) and impaired viral clearance by day 7 (Fig. 2G). This increased viral burden was accompanied by markedly elevated inflammatory cytokine levels in the lungs (Fig. 2H).

Together, these results demonstrate that HDAC1 and HDAC2 are essential for AM-mediated surfactant clearance and antiviral defense, highlighting their critical role in pulmonary immunity.

### HDAC1 and HDAC2 are required for early development and survival of Ams

AMs mainly originate from fetal monocytes that colonize the lung before birth and subsequently acquire a mature phenotype. To confirm whether HDAC1 and HDAC2 are required for the early establishment of AMs, we analyzed lungs from *Itgax*^Cre^*-*DKO mice at multiple postnatal time points (from day 3 to day 21). We observed a progressive increase in AM numbers and maturation status in control mice, with robust Siglec-F expression emerging between postnatal days 7 and 14 (Fig. 3A,B). In contrast, AMs from *Itgax*^Cre^*-*DKO mice exhibited markedly reduced Siglec-F expression (Fig. 3A) and significantly lower cell numbers compared with controls at each time point (Fig. 3B). To determine whether developmental proliferation was impaired, we assessed Ki-67 expression within AMs. Comparable frequencies of Ki-67⁺ AMs were detected in both groups (Fig. 3C,D), indicating that *Hdac1* and *Hdac2* deficiency may not significantly impair AM proliferation. Instead, these findings indicate that HDAC1 and HDAC2 may be more likely due to defects in the generation and survival of early AMs.

**Figure 3.**
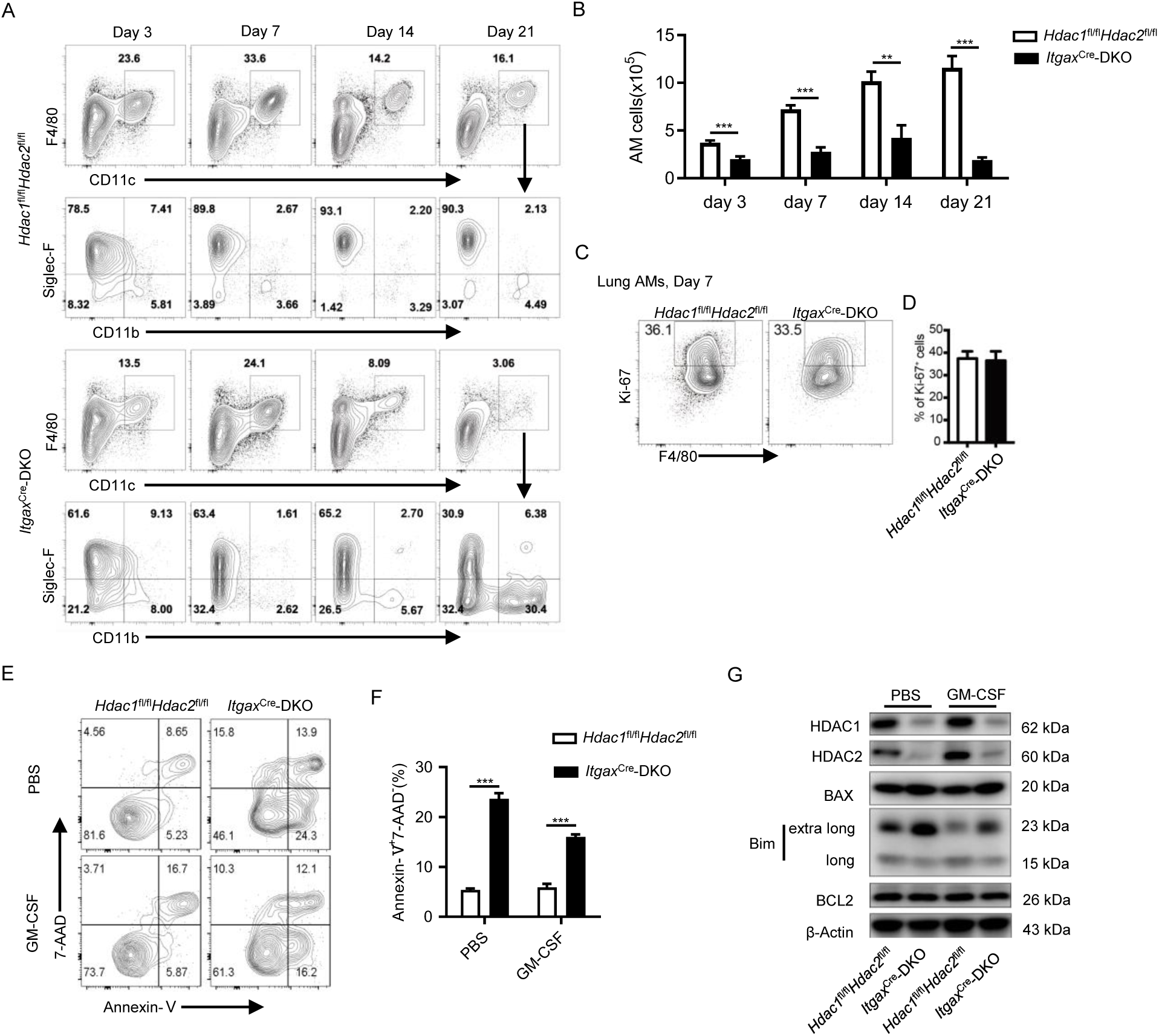
*Hdac1* and *Hdac2* are required for early development and maintenance of AMs. (A, B). FACS analysis of AMs in the lungs at indicated time points (A) and cell numbers (B) of AMs (CD11c^+^F4/80^+^) shown in (A) (n=5-16, pooled from two to four experiments). (C, D). FACS analysis of Ki-67^+^ AMs in the lungs of 7-day-old mice (C) and quantification of the cell percentages (D) (n=4). Data were representative of two independent experiments. (E, F). Apoptosis of AMs was analyzed by Annexin V and 7-AAD staining. BALF AMs were isolated by using CD11c microbeads and cultured *in vitro* for 24 hours with or without GM-CSF (20 ng/mL). Data were shown as mean ± SD from three technical repeats and were representative of two independent experiments. (G) Western blot analysis of apoptosis-related proteins in AM protein extracts. AMs were isolated as described in (E) and treated with GM-CSF or PBS for 30 minutes. Data were representative of two independent experiments. Data were shown as mean ± SEM. Statistical significance was determined by two-tailed unpaired Student’s *t* test. *, *p* < 0.05; **, *p* < 0.01; ***, *p* < 0.001.

To determine whether HDAC1 and HDAC2 deficiency affects AM survival, we next assessed the apoptosis rate of *Hdac1/2*-deficient AMs. At postnatal day 14, Western blot analysis confirmed efficient depletion of HDAC1 and HDAC2 in AMs isolated from *Itgax*^Cre^*-*DKO mice, confirming efficient Cre-mediated deletion (Fig. 3G). *In vitro* culture assays revealed increased apoptosis in AMs isolated from *Itgax*^Cre^*-*DKO mice (Fig. 3E, F). We then examined key signaling pathways known to regulate AM identity and survival, including GM-CSF receptor signaling, PPAR-γ, mTOR, and STAT5. Expression and activation of these regulators were comparable between control and *Itgax*^Cre^*-*DKO AMs (Fig. S4A-D), suggesting that canonical pathways required for AM differentiation and maintenance remain intact.

However, analysis of apoptotic regulators revealed increased expression of pro-apoptotic proteins, particularly Bim, whereas the anti-apoptotic factor Bcl-2 remained unchanged. RNA-seq analysis further revealed enrichment of apoptosis-related pathways among the differentially expressed genes in DKO AMs (Fig. S5). Therefore, HDAC1 and HDAC2 are indispensable for maintaining AM survival during early development by restricting apoptosis.

### The deacetylase activity of HDAC1 and HDAC2 is essential for maintaining AM survival

To determine whether their enzymatic activity is required for AM survival, we used a catalytically inactive HDAC1 mutant (H141A). Using an established culture system for generating AM-like cells, we overexpressed either wild-type HDAC1 or H141A mutant in AM-like cells derived from *Hdac1* and *Hdac2* double-deficient mice.

Unlike cells that overexpressed wild-type HDAC1, cells overexpressing the H141A mutant failed to restore the AM-like cell population in culture (Fig. 4A,B), indicating that the deacetylase activity of HDAC1 and HDAC2 is essential for maining AM survival. To further validate this requirement, we treated AMs *in vitro* with FK228, a selective inhibitor of HDAC1 and HDAC2. Consistently, inhibition of HDAC1 and HDAC2 enzymatic activity resulted in a marked increase in Bim both at the mRNA level (Fig. 4C) and protein level (Fig. 4D), and a higher proportion of apoptotic AMs (Fig. 4E-F). Importantly, knockdown of *Bim* in AMs partially rescued cell survival under these conditions (Fig. 4E, F), supporting a causal role of Bim in mediating apoptosis upon HDAC1 and HDAC2 inhibition.

**Figure 4.**
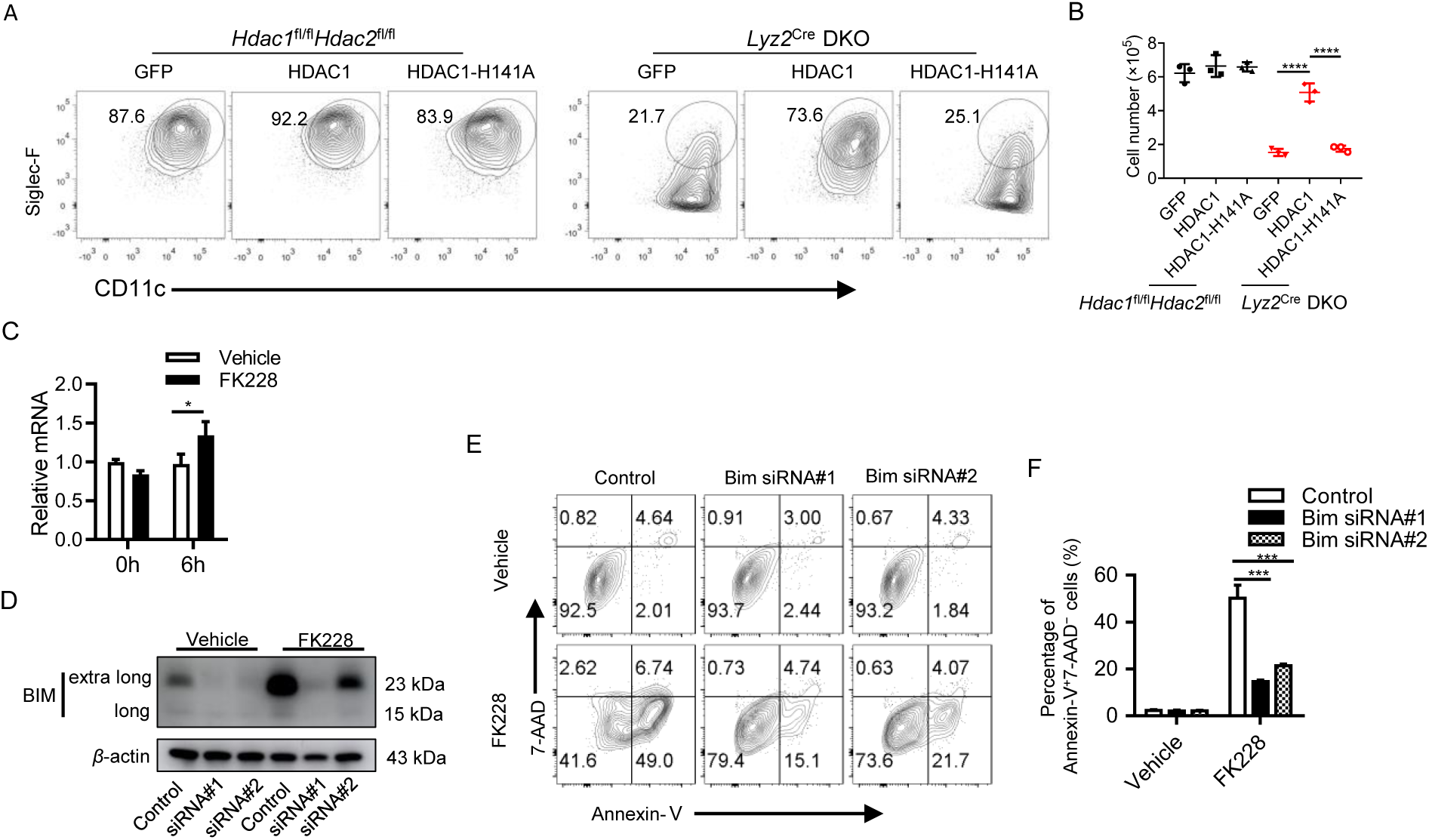
The deacetylase activity of HDAC1 and HDAC2 is essential for preventing AM apoptosis. (A, B). Representative flow cytometry analysis (A) and quantification (B) of AM-like cells derived from bone marrow Lin⁻ cells isolated from *Hdac1*^fl/fl^*Hdac2*^fl/fl^ and *Lyz2*^Cre^-DKO mice and transduced with GFP (control), HDAC1 or catalytically inactive HDAC1-H141A. Data were pooled from two independent experiments and shown as mean ± SD. Statistical significance was determined by one-way ANOVA with Sidak multiple comparisons test. ∗∗∗∗*p* < 0.0001. (C). Relative *Bim* mRNA level in BALF AMs, normalized to *Actb*. (D). Western blot analysis of BIM in protein extracts of BALF AMs with or without treatment of FK228 (50 nM). (E, F). Apoptosis of BALF AMs was analyzed by Annexin-V and 7-AAD staining (E) and percentage (F) the early apoptosis cells (Annexin-V^+^ 7-AAD^−^). The preparation of AMs was the same with that in (D). Data are shown as mean ± SD of three technical replicates and statistical significance was determined by an unpaired two-tailed Student’s *t* test. *, *p* < 0.05; **, *p* < 0.01; ***, *p* < 0.001.

Collectively, these data demonstrate that HDAC1 and HDAC2 maintain AM survival by repressing the pro-apoptotic gene *Bim* in a deacetylase-dependent manner.

### HDAC1 and HDAC2 regulate AM differentiation by interacting with LSD1

Given that *Bim* knockdown only partially rescued apoptosis in *Hdac1/2*-deficient AM-like cells, we next tested whether it could also rescue differentiation defects. Using an AM-like culture system with *Bim*-specific shRNA knockdown, we observed no significant improvement in AM differentiation (Fig. S6), indicating that HDAC1 and HDAC2 regulate AM differentiation through additional mechanisms.

To further investigate how HDAC1 and HDAC2 regulate AM differentiation, we performed co-immunoprecipitation (co-IP) of HDAC1 and HDAC2 in AM-like cells, followed by proteomic analysis. This analysis identified 40 shared interacting proteins between HDAC1 and HDAC2 (Fig. 5A), including subunits of the NuRD complex (MTA1/2/3, MBD2, RBBP4/7, GATAD2A/B), CoREST components (RCOR1/3), and the histone demethylase LSD1 (KDM1A), which is functionally cooperates with HDAC1 and HDAC2. Reciprocal endogenous co-IP further validated the interactions among HDAC1, HDAC2, and LSD1 (Fig. 5B), suggesting a coordinated role of this complex in regulating AM differentiation.

**Figure 5.**
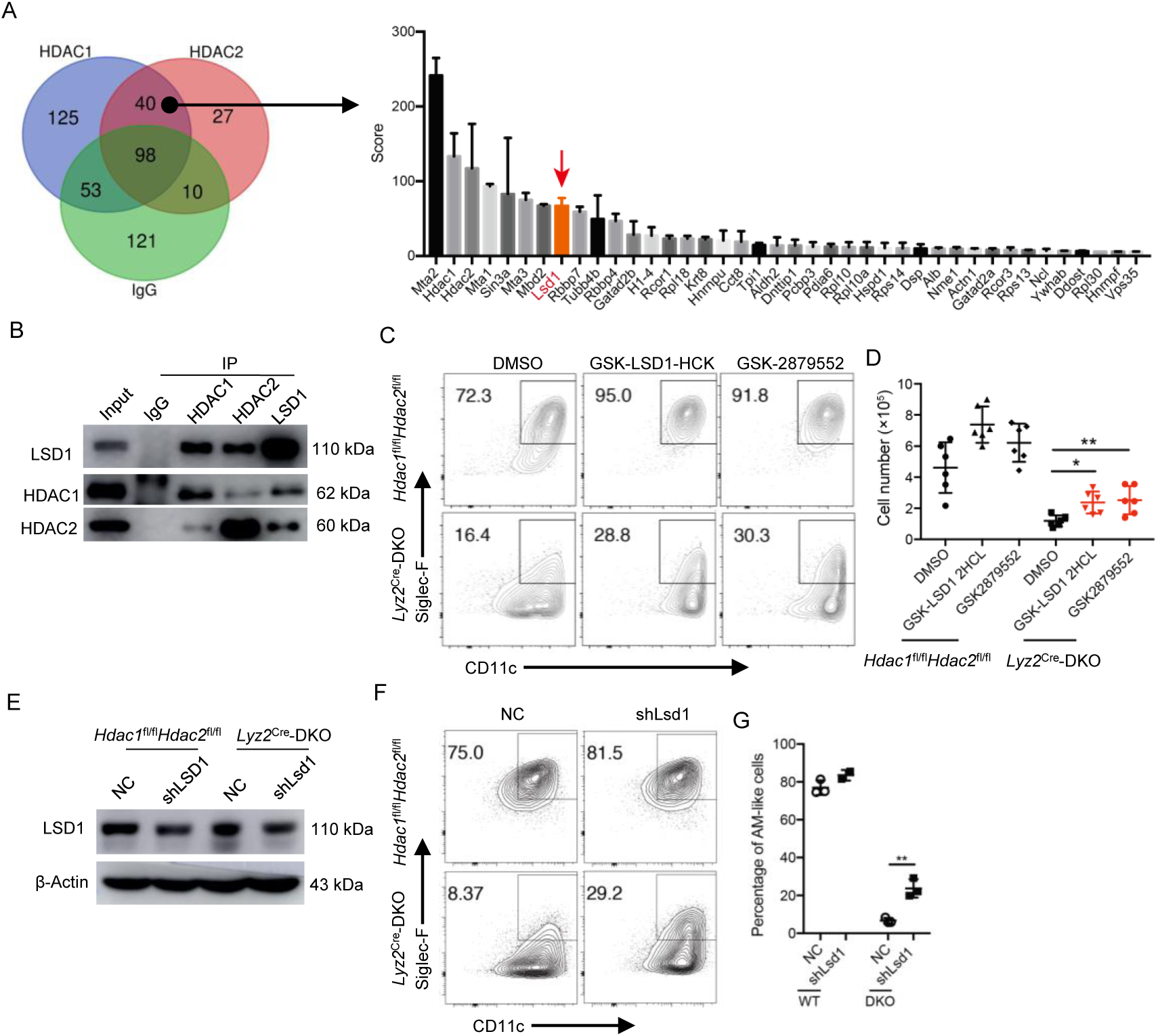
HDAC1/2 regulate the differentiation of AMs partially through LSD1. (A). Co-IP performed in AM-like cells using anti-HDAC1, anti-HDAC2, or control IgG antibodies, followed by proteomic analysis to identify interacting proteins. Left: Venn diagram of unique and shared proteins. Right: Top 40 proteins co-bound by HDAC1/2 ranked by enrichment scores. (B). Endogenous co-IP in AM-like cells using anti-HDAC1, anti-HDAC2, anti-LSD1, or IgG antibodies, followed by Western blot analysis. (C, D). Flow cytometry plots (C) and number (D) of AM-like cells derived from *Hdac1*^fl/fl^*Hdac2*^fl/fl^ and *Lyz2*^Cre^-DKO bone marrow treated with LSD1 inhibitors GSK-LSD1-HCl (50 μM) or GSK-2879552 (5 μM). (E). Western blot validation of LSD1 protein levels in AM-like cells transduced with shRNA targeting *Kdm1a* (shLsd1). (F and G). Flow cytometry analysis (F) and cell number (G) of AM-like cells derived from *Hdac1*^fl/fl^*Hdac2*^fl/fl^ and *Lyz2*^Cre^-DKO bone marrow Lin^−^ cells expressing shLsd1. Data were representative of two independent experiments (B and E) or three independent experiments (C, F and G), or pooled from two independent experiments (D, mean ± SD). Statistical significance was determined by one-way ANOVA followed by Sidak multiple Comparisons test (D) or by a two-tailed unpaired Student’s *t* test (G). ∗*p* < 0.05, ∗∗*p* < 0.01.

Notably, pharmacological inhibition of LSD1 in *Hdac1/2*-deficient cells enhanced AM-like differentiation and partially rescued differentiation defects (Fig. 5C, D). Similarly, shRNA-mediated knockdown of *Lsd1* (Fig. 5E) also improved differentiation in *Hdac1/2*-deficient cells (Fig. 5F, G).

Collectively, these data suggest that HDAC1/2 deficiency impaires AM differentiation, likely by disrupting the integrity of the HDAC-LSD1 complex and its downstream transcriptional programs.

## Discussion

The lung is constantly exposed to inhaled microbes and pollutants, with AMs serving as essential effectors for host defense, surfactant clearance, and lung development. In this study, we show that *Hdac1* and *Hdac2* double-deficiency led to severe AM defects, pulmonary alveolar proteinosis, and dramatically increased mortality after influenza infection. We further uncover that HDAC1 and HDAC2 maintained AM survival by repressing the pro-apoptotic protein Bim and promoted AM differentiation through interaction with LSD1. These findings define HDAC1 and HDAC2 as critical epigenetic regulators of AM homeostasis.

Epigenetic mechanisms, including histone modifications, DNA methylation, and noncoding RNAs, are central to AM development and function, modulating cellular responses to both developmental cues and environmental stimuli^35^. One well-established epigenetic mechanism is PRC2-mediated H3K27 trimethylation (H3K27me3). Loss of the PRC2 component SUZ12 impairs AM self-renewal and dramatically reduces AM numbers, highlighting the importance of histone methylation in AM biology^36^. Histone deacetylases (HDACs) constitute another major class of epigenetic regulators, among which HDAC3 has been shown to be essential for AM embryonic development, postnatal homeostasis, and maturation^37^. Yao et al. demonstrated that HDAC3 binds *Pparg* enhancers, and HDAC3 deficiency leads to mitochondrial oxidative dysfunction, impaired AM maturation, and disrupted AM homeostasis^37^. Our unpublished work also suggests that HDAC3 promoted AM proliferation via GM-CSF signaling, and maintained lipid metabolism through CPT1A, highlighting the critical role of HDAC3 in coordinating metabolic and developmental programs in AMs.

HDAC1, HDAC2, and HDAC3 all belong to class I HDACs, yet the roles of HDAC1 and HDAC2 in AM biology remain poorly defined. Despite their high sequence similarity (83%) and redundant functions in other cell types^23,26,38,39^, our data demonstrate that HDAC1 and HDAC2 are indispensable for AM survival and pulmonary homeostasis. Particularly, HDAC1, HDAC2, and HDAC3 regulate AM homeostasis by distinct mechanisms, indicating non-redundant functions within class I HDACs. HDAC3 drives AM development and maturation via PPARγ signaling^37^. However, in this study we showed that deficiency of HDAC1 and HDAC2 did not affect GM-CSFR expression, PPARγ activation, or STAT5 signaling, but instead primarily triggered extensive apoptosis during early AM establishment, accompanied by Bim upregulation and widespread transcriptomic changes enriched in apoptosis pathways. HDAC1 and HDAC2 act as catalytic cores within co-repressor complexes such as Sin3^40^, NuRD^41^, and CoREST^42^, which likely confer distinct transcriptional repression programs compared with HDAC3-associated complexes.

Different epigenetic mechanisms often cooperate to fine-tune gene transcription. For instance, HDAC-mediated deacetylation and LSD1-mediated demethylation can act synergistically within shared co-repressor complexes to define cell fate decisions and maintain cellular fitness in response to diverse stimuli^43,44^. In this study, we validated a physical interaction between HDAC1/2 and LSD1. Moreover, both pharmacological inhibition and genetic knockdown of LSD1 partially rescued the AM defects resulting from HDAC1 and HDAC2 deficiency. These findings support a model in which HDAC1/2 and LSD1 function cooperatively to guide AM development and homeostasis. We propose that, in the absence of HDAC1 and HDAC2, the HDAC-LSD1 complex becomes destabilized, leading to dysregulated transcriptional programs at LSD1 target loci required for AM development, ultimately resulting in AM loss.

## Method

### Mice

C57BL/6J (Jax 000664) and CD45.1 (Jax 002014) were purchased from the Jackson Laboratory. *Lyz2*-Cre mice (Jax 004781) were purchased from Model Animal Research Center of Nanjing University, China. *Itgax*-Cre mice (Jax 007567) were a gift from Dr. Wanli Liu (Tsinghua University, China). *Hdac1*^fl/fl^ mice^23^ on the B6 background were kindly provided by Dr. Fanglin Sun (Tongji University, China). *Hdac2*^fl/fl^ mice^45^ on the B6 background were kindly provided by Dr. Jisong Guan (Tsinghua University, China). *Hdac1* and/or *Hdac2* cKO mice were generated by crossing *Hdac1*^fl/fl^ and *Hdac2*^fl/fl^ mice with *Itgax*-Cre or *Lyz2*-Cre mice. The ROSA26-loxP-Stop-loxP RFP reporter mice^33^ were a gift from Dr. Yuncai Liu (Tsinghua University, China). All mice were used at 8 to 10 weeks of age unless otherwise stated. Mice were maintained under specific pathogen-free conditions in the animal facility of Tsinghua University. All animal procedures were approved by the Institutional Animal Care and Use Committee at Tsinghua University.

### Cells Isolation

Lung cells were obtained as described previously with slight modifications^46^. Briefly, for adult mice, 3 mL HBSS were injected into the right ventricle to flush blood from the lungs. The lungs were removed and minced, then digested in 5 mL HBSS containing 5% FBS, collagenase type 1A 0.5 mg/ml (Sigma, C9891-1G) and 0.05 mg/mL DNase I (Roche Molecular Biochemicals, Indianapolis, IN, USA) for 1 hour at 37℃, and the samples were vortexed every 15 min to resuspend tissue fragments. For the mice younger than 3 weeks old, lung perfusion step was skipped. The volume of digestion solution was 2.5 mL, and the time for incubation was 45 min. The digested content was passed through a 70 μm cell strainer to obtain a homogeneous cell suspension.

Cell suspension of small intestine was obtained as described previously ^47^. Briefly, the small intestine was removed and carefully cleaned of the mesentery, Peyer’s patches, fat and content. The epithelial cells were removed by two sequential incubations in HBSS with 5% FCS and 2 mM EDTA at 37℃ for 20 min. The tissue fragments were minced and incubated in RPMI 1640 with 5% FCS, 10 U/mL collagenase CLSPA (Worthington Biochemical), and 0.05 mg/mL DNase I at 37℃ under agitation for 40 min. Cell suspensions were passed through a 100-μm strainer. The cells were further enriched with 40% Percoll (GE Healthcare) by centrifugation at 2000g for 10 min.

Peritoneal lymphocytes were obtained by lavage with two washes of 5 mL PBS with 3% FCS. Splenic lymphocytes were collected by mashing spleen through 70 μm cell strainer. BM cells were harvested from femur and tibia by flushing the marrow out with PBS with 3% FCS.

### Flow Cytometry

Cells were incubated with in-house anti-CD16/32 antibody or rat IgG (Jackson Laboratories, USA) for 10 min to block Fc receptor binding. Cells were then stained with indicated antibodies for 30 min at 4 °C for cell surface marker analysis. For Ki-67 staining, surface markers were first stained, followed by fixed and intracellular staining using the Foxp3/Transcription Factor Staining Buffer Set kit (eBioscience) according to manufacturer’s instructions. For apoptosis analysis, cells were stained using tthe AnnexinⅤapoptosis detection kit (Biolegend) according to manufacturer’s instructions.

The following antibodies were used for cell staining: fluorochrome conjugated monoclonal antibodies (mAbs) against mouse antigens: CD11c (clone N418); CD45 (clone 30-F11); F4/80 (clone BM8); CD11b (clone M1/70); Siglec-F (clone E50-2440); CD103 (clone 2E7); CD24 (clone M1/69); CD64 (clone X54-5/7.1); MHCII (clone M5/114.15.2); Ly6G (clone 1A8); CD3e (clone 145-2c11); CD8α (clone53-6.7); CD4(clone GK1.5); CD45R (clone RA3-6B2); CD19 (clone eBio1D3); Ly6C (clone H1.4); CD45.2 (clone 104); CD45.1(clone A20); Ki-67 (clone SolA15); corresponding isotype controls; Viability dye 7-AAD was used to discriminate dead cells. All these antibodies and reagents were purchased from eBioscience, BD Biosciences or Biolegend. GM-CSFR α-PE was purchased from R&D.

LSRⅡ flow cytometers (Becton Dickinson) were used for cell analysis; FACSAria Ⅲ instruments (Becton Dickinson) were used for cell sorting. FACS data were analyzed and displayed with the FlowJo software (Tree Star, Ashland, OR, USA).

### Quantitative real-time PCR (RT-PCR)

Total RNA from tissues or isolated cells was extracted by TRIzol reagent (Invitrogen, Grand Island, NY, USA) according to the manufacturer’s instructions. cDNA was prepared by reverse transcription with the PrimeScript RT Reagent Kit (Takara, Shiga, Japan). Amplification was performed on an ABI 7900 Real-Time PCR System (Applied Biosystems, Grand Island, NY, USA) with the SYBR PrimeScript RT-PCR Kit (Takara, Shiga, Japan). Relative gene expression was calculated using the comparative method for relative quantitation by normalization to the housekeeping gene *Actb* or *Hprt*. Primers used: *Actb* (forward)5’-ATGCTCCCCGGGCTGTAT-3’, (reverse)5’-CATAGGAGTCCTTCTGACCCATTC-3’, *Hprt*:(forward)5’-TAATGTA ATCCAGCAGGTCAG-3’, (reverse)5’-TCATGGACTGATTATGGACAG-3’, *Hdac1*:(forward)5’-GATATTCACCATGGCGATG-3’, (reverse)5’-CCCAGTTCCTG GGAAGTAC-3’, *Hdac2*:(forward)5’-AACCGGCAACAAACTGAT-3’, (reverse)5’-ATGGCAAGCACAATATCATT-3’, *Sp-a*:(forward)5’-CGGCTCTGGTACACATCTC T-3’, (reverse)5’-TCTGCAAACAATGGGAGTC-3’, *Sp-d*:(forward)5’-TCCTGGAG GTCCACTTAGTC-3’, (reverse)5’-GGTTTGCCAGGACCTATG-3’, *Il1b*:(forward)5’-GCTTGGGATCCACACTCTC-3’, (reverse)5’-TGATATTCTCCAT GAGCTTTGT-3’, *Ccl2*:(forward)5’-GCTGGTGATCCTCTTGTAGC-3’, (reverse)5’-GCTCAGCCAGATGCAGTT-3’, *Tnf*:(forward)5’-CCACTTGGTGGTTTGCTAC-3’, (reverse)5’-CCCAAAGGGATGAGAAGTT-3’, *Il6*:(forward)5’-TCCAGAAGACCA GAGGAAAT-3’, (reverse)5’-AGTTGTGCAATGGCAATTC-3’, *Flu virus*: (forward) 5’-GGACTGCAGCGTAGACGCTT-3’, (reverse)5’-CATCCTGTTGTATATGAGGC CCAT-3’.

### Bronchoalveolar lavage fluid (BALF) collection and analysis

The trachea was exposed and cannulated. For adult mice, the lungs were flushed three times with 0.6 ml PBS. BALF samples were centrifuged to separate cells from supernatants. The supernatants were stored at −80°C. Red blood cells in BALF were lysed with RCRB buffer. Leukocytes were washed once to obtain single-cell suspensions for experiments. For protein concentration quantification, BALF supernatants were measured by BCA Protein Assay Kit (Thermo Scientific).

### Immunoblot analysis

Cell lysates were separated by 4-20% gradient SDS-PAGE (Beyotime, China) and transferred to 0.45- or 0.22-μm Immobilon-P polyvinylidene difluoride membrane (Millipore) by electroblotting. The membranes were blocked with QuickBlock blocking buffer (Beyotime, China) prior to antibody incubation. Proteins were visualized with enhanced chemiluminescence ECL kit (YEASEN, China). Amersham Image 600 system (GE Healthcare) was used for the chemiluminescence signal detection. The following antibodies were used for incubation: SP-A (Santa Cruz, sc-13977); SP-D (Santa Cruz, sc-13980); β-Tubulin (EASYBIO, BE0025); GAPDH (Proteintech, HRP-60004); GM-CSFR β (Santa Cruz, sc-678); β-Actin (CST, 4967S); HDAC1 (CST, 34589S); HDAC2 (CST, 57156S); p-PPAR γ (Affinity, AF3675); PPAR-γ (CST, 2435T); p-STAT5 (CST, 4322S); STAT-5 (CST, 94205T); p-mTOR (CST, 5536T); mTOR (CST, 2983S); β-Actin (EASYBIO, BE0022); BAX (Selleck, A5131); Bim (CST, 2819T); BCL-2 (Affinity, AF6139); goat anti-rabbit IgG-HRP conjugated antibody (ZSGB-BIO, ZB-5301); goat anti-mouse IgG-HRP antibody (ZSGB-BIO, ZB-2305).

### Histopathological analysis of the lung

The lungs were removed and fixed with 4% paraformaldehyde for at least 24 hours. The tissues were embedded in paraffin. Lung sections were stained with H&E (hematoxylin and eosin) or PAS (Periodic acid-Schiff) processed according to standard protocols.

### Influenza virus infection

Influenza A/Puerto Rico/8/1934 (H1N1) (PR8) virus stocks were kindly provided by Dr. Xu Tan (Tsinghua University, China). For influenza infection studies, 8-week-old sex-matched mice were anesthetized with isoflurane and intranasally infected with 500 TCID_50_ (50% tissue culture infective dose) of influenza PR8 virus in 30 µl PBS. Body weight and survival of mice were monitored daily for 12 days after infection. Mice were euthanized for experiments at indicated time points after infection. Viral load in the lungs was performed as described^48^. Briefly, whole lungs were harvested and homogenized to extract total RNA. qPCR was performed with primers specific for the matrix protein of the influenza virus. Viral RNA levels were normalized to *Actb*.

### Construction of BM chimeras

B6 CD45.1 recipients were lethally irradiated by X-ray irradiation (5.5 Gy, twice), and then intravenously transferred with a combination of 4×10^6^ BM cells from B6 CD45.1 mice and indicated donor BM cells mixed at a 1:1 ratio. Chimeras were given drinking water with antibiotics (1 mg/ml neomycin and 100 μg/ml polymyxin B, Inalco Pharmaceuticals) for 2 weeks. Chimeras were analyzed 8 weeks after reconstitution.

### In vitro GM-CSF stimulation of AMs

AMs were collected from BALF of 2-week-old mice and enriched using CD11c microbeads (Miltenyi) according to manufacturer’s instructions. For apoptosis analysis, enriched AMs were cultured in complete DMEM medium with 20 ng/mL GM-CSF for 24 hours. Adherent AMs were detached by incubating with Accutase (sigma) for 20 min. For Western blot analysis, enriched AMs were rested in complete DMEM medium for 2 hours and then stimulated with 20 ng/mL GM-CSF for 30 min.

### Differentiation of AM-like cells

AM-like cells were generated as described previously. Briefly, total bone marrow (BM) cells were harvested and treated with RBC lysis buffer. Cells were then resuspended and cultured in complete low-glucose DMEM supplemented with 20 ng/ml GM-CSF and 2 ng/ml TGF-β. After 7 days of culture, the medium was replaced with complete low-glucose DMEM containing 20 ng/ml GM-CSF, 2 ng/ml TGF-β, and 100 nM Rosiglitazone for an additional 2 days. On day 9 of differentiation, non-adherent cells were discarded, and adherent macrophages were maintained as AM-like cells.

### RNA sequencing (RNA-seq) and data analysis

AMs (CD11c^+^F4/80^+^) were sorted from lungs of 2-week-old mice. Total RNA was extracted by TRIzol reagent according to the manufacturer’s instructions. Library construction and data processing were performed by Beijing Genomics Institute. Libraries were sequenced on an Illumina HiSeq 4000 platfform using a single-end 50 bp reads (SE50) strategy. Reads were aligned and quantified, and gene expression levels were normalized as fragments per kilobase of exon per million mapped reads (FPKM). RNA-seq experiments were performed with three biological replicates per group. Differentially expressed genes (DEGs) were identified using DESeq2 with a cutoff of fold change ≥1.8 and adjusted *p* < 0.05. Gene set enrichment analysis (GSEA) was performed using clusterProfiler with Gene Ontology (GO) gene sets and the GSKB database. The RNA-seq data have been deposited in the Gene Expression Omnibus (GEO) under accession number GSE155965.

### RNA interference (RNAi)

Small interfering RNA (siRNA) targeting mouse *Bim* and non-targeting control siRNA were obtained from GenePharma (Suzhou, China). The sequence of Bim siRNA#1 and siRNA#2 were GACGAGUUCAACGAAACUUACTT and AUUGUCCACCUUCU CUGUCACTT, respectively. A total of 4×10^4^ BALF AMs were rested in complete DMEM supplemented with 20 ng/ml GM-CSF for 24 h. siRNAs were transfected into AMs using TransIT-TKO transfection reagent (Mirus Bio) according to the manufacturer’s instructions. Cells were used for FK228 treatment or protein extraction at 48 h post-transfection.

### Retroviral transduction

As described previously, lineage-negative (Lin[) BM cells from Cas9 knock-in mice were enriched via negative magnetic selection using an antibody cocktail including anti-TER119, anti-CD2, anti-CD3, anti-CD45R, anti-CD8, anti-CD11b, and anti-Ly6G. Retroviruses were produced in Plat-E cells by transfection with the following plasmids: pRVKM-U6-NC, pRVKM-U6-Lsd1 shRNA, pRVKM-U6-Bim shRNA-CMV-mAmetrine, pMYS-IRES-EGFP, pMYS-HDAC1-IRES-EGFP, or pMYS-HDAC1(H141A)-IRES-EGFP plasmids.

Lin⁻ cells were prestimulated overnight with 100 ng/ml stem cell factor (SCF), 10 ng/ml IL-6, and 6 ng/ml IL-3, and then transduced on Retronectin-coated plates (T100B; Takara). At 48 h post-transduction, mAmetrine⁺ or EGFP⁺ cells were sorted by flow cytometry and cultured for 7 days in medium containing 20 ng/ml GM-CSF and 2 ng/ml TGF-β, followed by an additional 2 days in medium supplemented with 20 ng/ml GM-CSF, 2 ng/ml TGF-β, and 100 nM rosiglitazone.

### Co-Immunoprecipitation and mass spectrometry analysis

AM-like cells prepared in three 10-cm dishes were lysed with 1 mL of lysis buffer per dish. Lysates were incubated overnight with control IgG (CST, 2729s), anti-HDAC1 (CST, 34589S), or anti-HDAC2 (CST, 57156S) antibodies. Protein A/G magnetic beads (Millipore, 16-663) were added the next day and incubated for 2 h. Bead complexes were magnetically isolated and washed three times with lysis buffer, followed by resuspension in 1× SDS loading buffer. Samples were resolved by SDS-PAGE and all protein bands were excised from the gel and subjected to mass spectrometry analysis at the Technology Center for Protein Sciences, Tsinghua University.

### Statistical analysis

Statistical analyses were performed with GraphPad Prism 7.0 (GraphPad Software). Data were analyzed by ANOVA followed by Bonferroni multiple comparison test or by unpaired, two-tailed Student’s t test. Data were presented as mean ± SEM or mean ± SD, and p <0.05 (*) was considered statistically significant.

## AUTHOR CONTRIBUTIONS

L.W. and W.L. contributed to study design. W.L., M.L., and T.W. performed experiments and analyzed the data. M.L. and Z.H. helped with AM culture and Western blot analysis. Y.W. and X.L. contributed to animal experiments and RNA-seq. L.W., W.L., T.W. and M.L. contributed to manuscript preparation. The study was supervised by L.W.

## ACKNOWLEDGMENTS

We thank members of the Wu laboratory for their help. This study was supported by National Natural Science Foundation of China (82388101 to L.Wu), China Postdoctoral Science Foundation (2024M762066 to M. Luo) and Postdoctoral Fellowship Program of CPSF (GZC20231626 to M. Luo). We would like to thank Dr. Fanglin Sun, Dr. Wanli Liu, Dr. Jisong Guan, Dr. Yuncai Liu and Dr. Wen Zhang for providing the mice. We would like to thank Dr. Xu Tan and Dr. Xiuxiu Yang for providing the influenza virus and experimental guidance.

---

**Figure S1.**
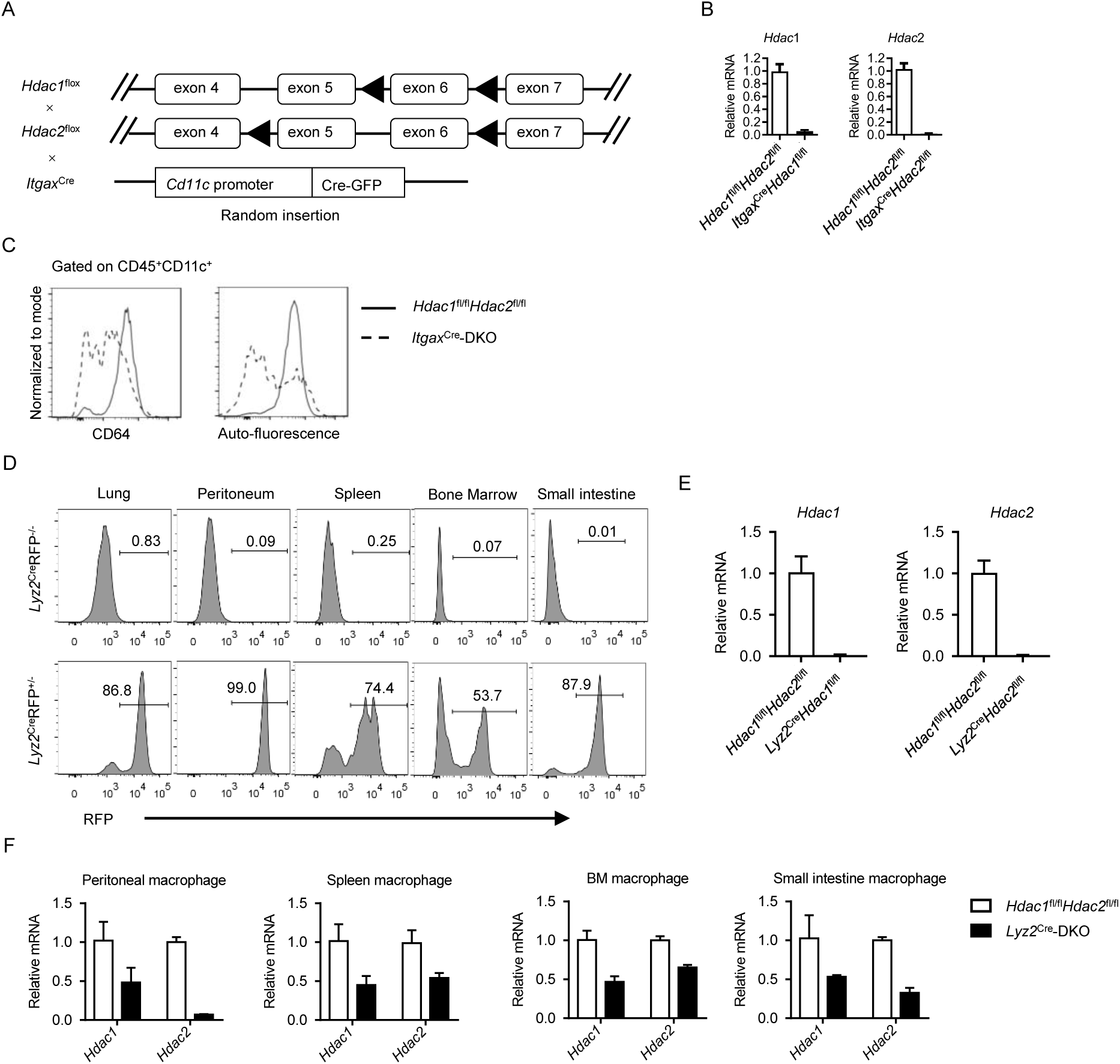
*Itgax*^cre^ and *Lyz2*^cre^ efficiently delete *Hdac1* and *Hdac2* in AMs. (A). Schematic diagram of the used mice model. (B). Relative mRNA level of *Hdac1* and *Hdac2* in AMs from indicated mice, normalized to *Actb* expression. (C). Expression of CD64 and autofluorescence of cells gated as CD45^+^CD11c^+^ cells (n=3). Data were representative of two independent experiments. (D). Representative frequency of RFP^+^ cells in macrophages isolated from lung (AM), spleen, bone marrow (BM), peritoneum and small intestine. (E). Relative mRNA level of *Hdac1* and *Hdac2* in isolated AMs from indicated mice, normalized to *Actb* expression. (F). Relative mRNA level of *Hdac1* and *Hdac2* in macrophages isolated from peritoneal, spleen, BM and small intestine from indicated mice, normalized to *Actb* expression. Data were shown as mean ± SEM, and were representative of two independent experiments.

**Figure S2.**
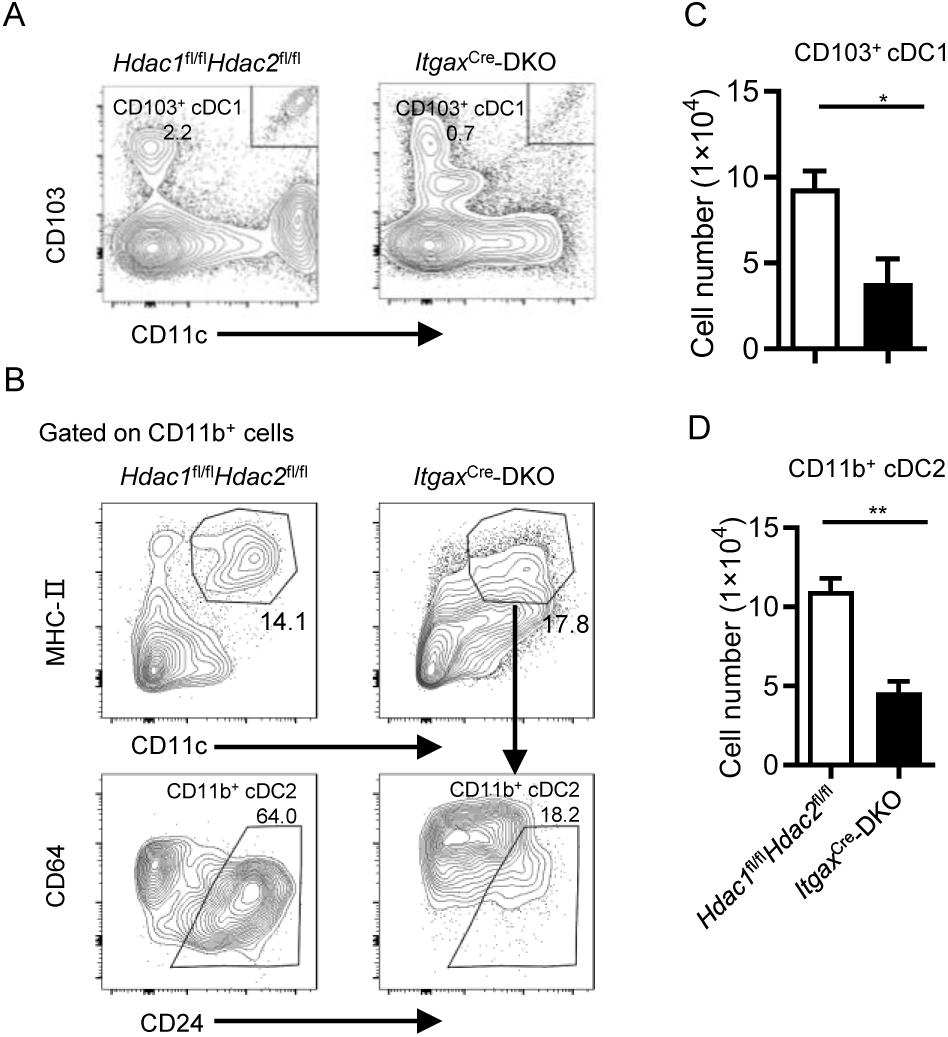
Dual loss of *Hdac1* and *Hdac2* impairs cDC homeostasis in the lung. (A, B). Representative flow cytometry plots of lung CD103^+^ cDC1 (A) and CD11b^+^ cDC2 (B). (C, D) Cell numbers of lung CD103^+^ cDC1 (C) and CD11b^+^ cDC2 (D) (n=3-5). Data were representative of three independent experiments, and shown as mean ± SEM. Statistical significance was determined by unpaired two-tailed Student’s *t* test. *, *p* < 0.05; **, *p* < 0.01; ***, *p* < 0.001.

**Figure S3.**
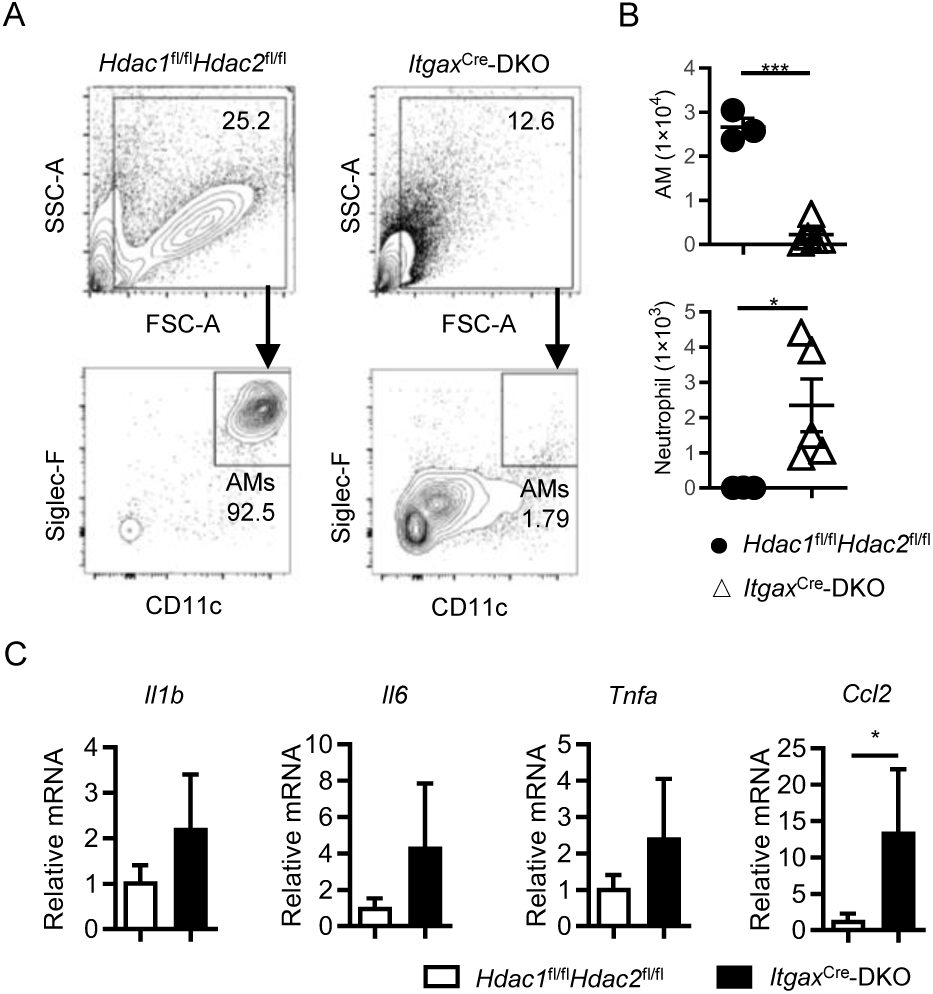
BALF AMs and neutrophils in *Hdac1* and *Hdac2* double deficient mice. (A, B). Representative flow cytometry plots (A) and cell numbers (B) of AMs and neutrophils (gated on 7-AAD⁻CD45^+^ population) in BALF (n=3-5). Data were representative of three independent experiments. (C). Relative mRNA levels of *Il1b, Il6, Tnf* and *Ccl2* in lung tissue, normalized to *Hprt* expression (n=3-4). Data were shown as mean ± SEM. Statistical significance was determined by unpaired two-tailed Student’s *t* test. *, *p* < 0.05; **, *p* < 0.001.

**Figure S4.**
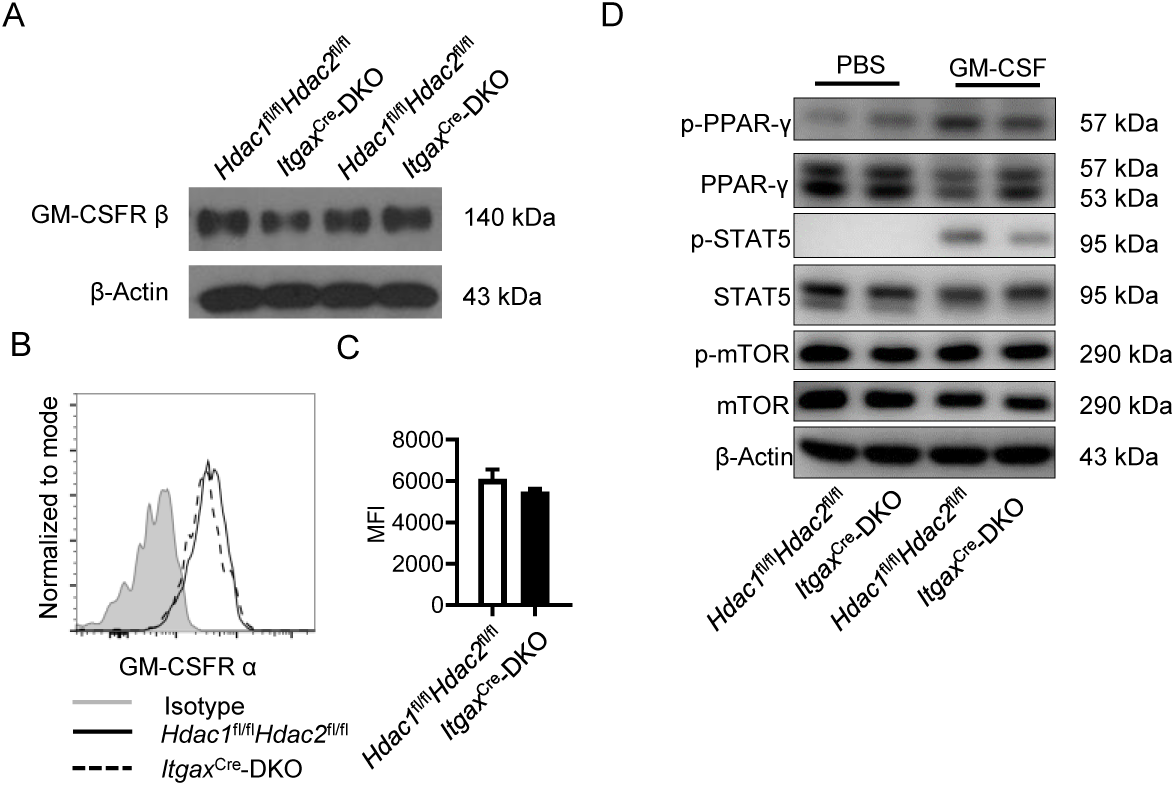
Dual loss of *Hdac1* and *Hdac2* in AMs shows no effect on GM-CSF signal transduction. (A). Western blot analysis of GM-CSFR β in AM protein extracts from 2-week-old mice. (B, C). Flow cytometric analysis of GM-CSFR α on AMs from 2-week-old mice (n=4-5). Data were shown as mean ± SD. Statistical significance was determined by two-tailed unpaired Student’s *t* test. (D). Western blot of HDAC1, HDAC2, total and phosphorylated PPAR-γ, STAT5, and mTOR in AM lysates from BALF. BALF AMs were isolated by using CD11c microbeads and cultured *in vitro* for 30 min with or without GM-CSF (20 ng/mL). Data were representative of two independent experiments.

**Figure S5.**
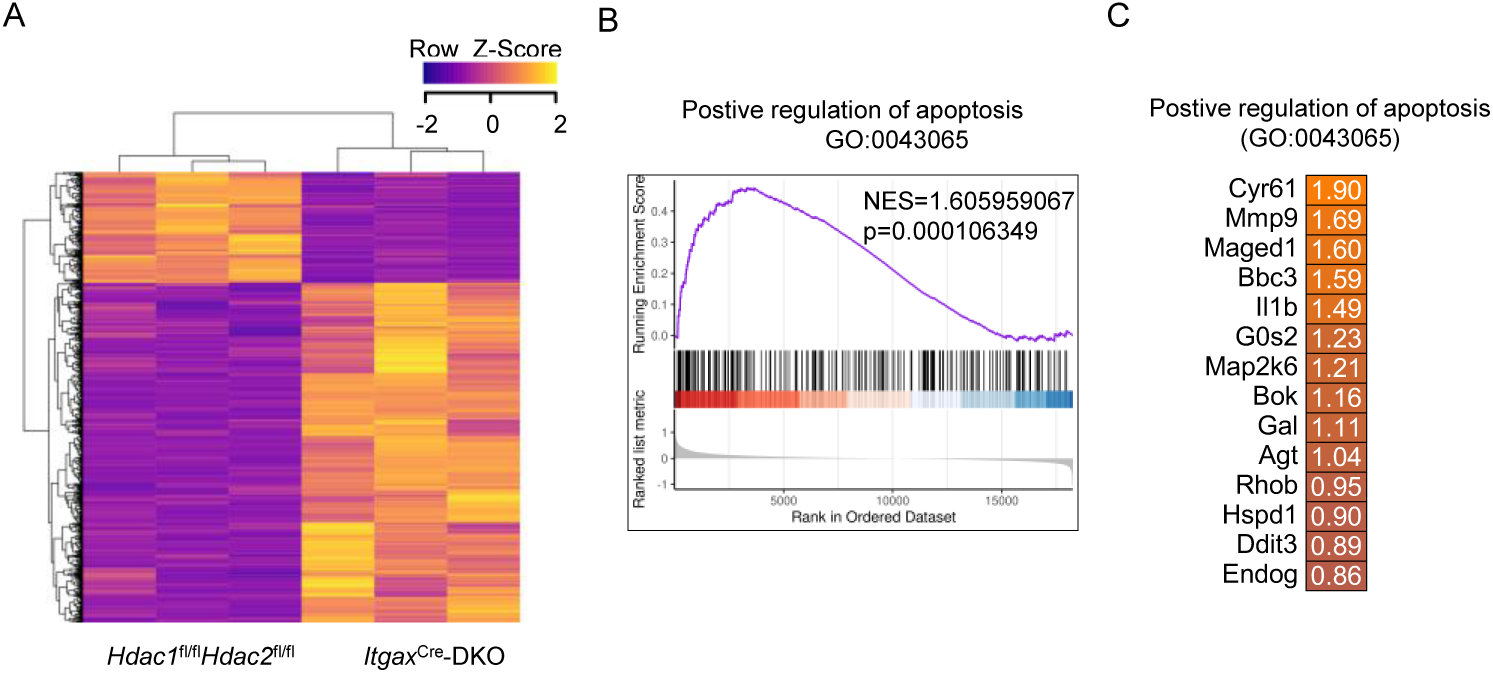
Dual loss of *Hdac1* and *Hdac2* enhances expression of apoptosis genes in AMs. (A). Heat map of differentially expressed genes in AMs sorted from the lungs of 2-week-old *Hdac1*^fl/fl^*Hdac2*^fl/fl^ and *Itgax*^Cre^-DKO mice (n=3 per group). Genes with a ≥1.8-fold change were considered differentially expressed. Expression values were normalized as Z-scores. (B) Gene Set Enrichment Analysis (GSEA) showing enrichment of the ‘positive regulation of apoptotic signaling pathway’ in *Hdac1/2*-deficient AMs. (C) Differentially expressed genes within the pathways shown in (B).

**Figure S6.**
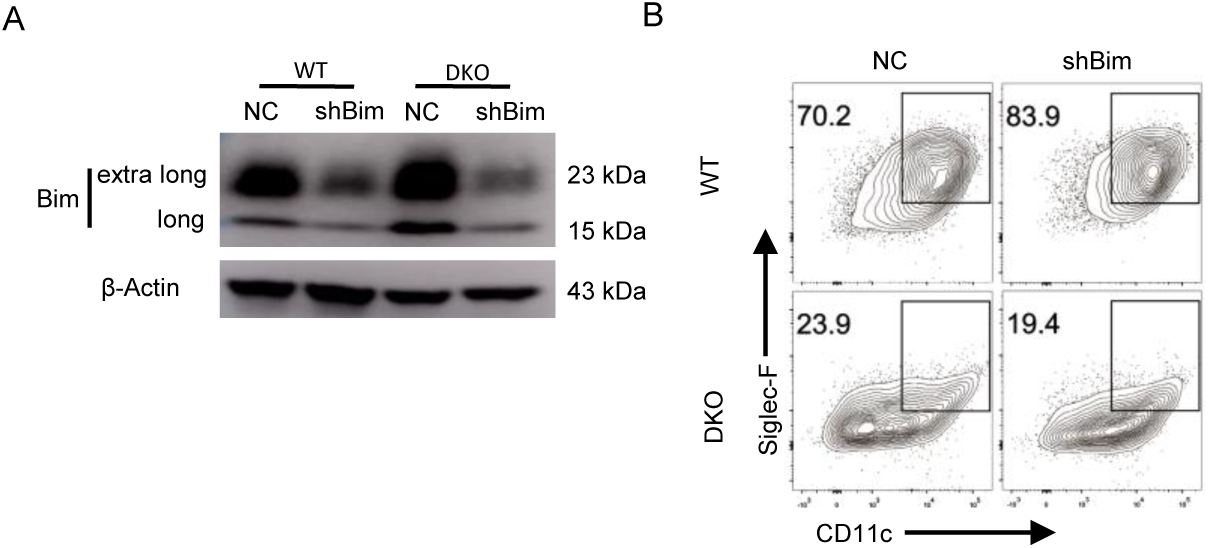
*Bim* knockdown does not rescue abnormal AM differentiation caused by HDAC1 and HDAC2 double-deficiency. (A). Western blot validation of BIM protein levels in AM-like cells transduced with *Bim*-targeting shRNA. (B). Flow cytometric analysis of AM-like cells derived from *Hdac1*^fl/fl^*Hdac2*^fl/fl^ and *Lyz2*^Cre^-DKO bone marrow Lin^−^ cells expressing shBim. Data were representative of three independent experiments.

